# Myeloperoxidase and Eosinophil Peroxidase Inhibit the *in vitro* Endotoxin Activities of Lipopolysaccharide (LPS) and Lipid A and Increase Survival in an *in vivo* Mouse LPS LD90 model

**DOI:** 10.1101/499855

**Authors:** Robert C. Allen, Mary L. Henery, John C. Allen, Roger J. Hawks, Jackson T. Stephens

**Affiliations:** Department of Pathology, Creighton University School of Medicine, Omaha, NE, USA; Exoxemis, Inc, Omaha, NE, USA; Concord Biosciences, LLC, Concord, OH, USA

## Abstract

Myeloperoxidase (MPO) and eosinophil peroxidase (EPO) are cationic leukocyte haloperoxidases with potent microbicidal and detoxifying activities. MPO selectively binds and kills specific Gram-positive bacteria (GPB) and all Gram-negative bacteria (GNB) tested. Endotoxin, i.e., lipopolysaccharide (LPS) comprising a toxic Lipid A component, is a characteristic of all GNB. The possibility that haloperoxidases bind to and inhibit the endotoxin of GBN was considered and tested by contacting MPO and EPO with LPS and Lipid A and measuring for inhibition of endotoxin activity using either the *in vitro* gel or chromogenic Limulus amebocyte lysate (LAL) assays. Contacting MPO and EPO with LPS purified from *Escherichia coli* O55:B5 and with diphosphoryl Lipid A purified from *E. coli* F583 inhibited their endotoxin activities in proportion to the natural log of the MPO or EPO concentration. Although MPO is less cationic than EPO, MPO consistently demonstrated inhibition of endotoxin activity that is about threefold superior to EPO. Haloperoxidase enzymatic activity was not required for inhibition, and MPO haloperoxidase action did not increase endotoxin inhibition. MPO and EPO inhibition of LPS endotoxin activity was also measured using a 90% lethal dose (LD90) mouse model studied over a five-day period. Based on Kaplan Meier survival analysis, MPO significantly increased mouse survival in a dose-dependent manner. EPO was less effective. In conclusion, contacting MPO and EPO with LPS and Lipid A inhibits *in vitro* endotoxin activities, but inhibition is independent of haloperoxidase enzymatic function. MPO significantly increases mouse survival against LPS in an *in vivo* LD90 endotoxin model.

## Introduction

Myeloperoxidase (MPO) is a unique dimeric heme A glycoprotein produced by neutrophil and monocyte leukocytes (1, 2). Eosinophil leukocytes produce a monomeric eosinophil peroxidase (EPO) with moderate homology to MPO (72.4% at the nucleotide and 69.8% at the amino acid level) (3, 4). Both MPO and EPO are cationic, but EPO is more cationic than MPO (5). Both enzymes are haloperoxidases; i.e., MPO and EPO catalyze the oxidation of chloride and bromide, respectively. Both enzymes have haloperoxidase activities that are highly microbicidal (6, 7). MPO production in neutrophils is abundant and influenced by the extent of stimulated myelopoietic turnover (8, 9).

MPO selectively binds and kills some specific gram-positive bacteria (GPB), but binds and kills all gram-negative bacteria (GNB) tested (10). In addition to microbial killing, the haloperoxidase activity of MPO has been reported to inactivate microbial toxins, including diphtheria toxin, tetanus toxin, and *Clostridium difficile* cytotoxin (11, 12). Like microbicidal action, the toxin-destroying activities of MPO require haloperoxidative enzymatic action that is hydrogen peroxide and halide-dependent. There are no reports of MPO binding or inhibition of the endotoxin activity of lipopolysaccharide (LPS) or Lipid A.

The cell envelope of GNB is composed of a cell wall with an inner cytoplasmic cell membrane and an outer cell wall membrane presenting LPS with its toxic Lipid A component (13). Release of endotoxin secondary to GNB lysis causes severe toxemia, i.e., septic shock. The MPO binding observed for all GNB tested (10, 11) suggests the possibility that endotoxin, a characteristic component of all GNB (14-16), might be responsible for such binding. The present studies were designed to determine if MPO binding to GNB might involve direct binding to endotoxin, and if such binding might inhibit the endotoxin activity of LPS and Lipid A.

The haloperoxidase activity of MPO is potent, and might be considered potentially toxic. Our contention is that MPO lethal action is binding-specific and focused, and that the highly reactive products of haloperoxidase action are temporally and physically restricted to the proximate site of enzyme binding (11). Recently, interesting evidence has been presented that blockade or genetic deletion of MPO increases mortality associated with LPS toxicity, and as such, MPO appears to protect against the adverse effects of endotoxin (17). The *in vitro* and *in vivo* research described herein provides direct empirical evidence for MPO and EPO contact inhibition of LPS and Lipid A endotoxin activities.

## Materials and Methods

### Enzymes

The haloperoxidases, porcine myeloperoxidase (MPO) and porcine eosinophil peroxidase (EPO), were produced by Exoxemis Inc. The porcine MPO used was 98.9% pure by ultraperformance liquid chromatography (RP-UPLC) and 100% pure by molecular size exclusion high performance liquid chromatography (SEC-HPLC). The MPO had a reinheitszahl (RZ; A_430nm_/A_280nm_) of 0.79. The porcine EPO used was 99.2% pure by reverse phase high performance liquid chromatography and had a Reinheitszahl (RZ; A_415nm_/A_280nm_) of 0.96. Glucose oxidase (GO) was isolated from *Aspergillus niger* and purified by Exoxemis Inc. Its final purity was 99.8% by RP-HPLC and 99.9% by SEC-HPLC.

### Endotoxins

Lipopolysaccharide (LPS) purified from *E. coli* O55:B5 was purchased from Sigma-Aldrich (L4524). The manufacturer specification for LPS endotoxin activity was 3 x 10^6^ endotoxin units (EU) per mg. Using standard LPS dilutions, our results were consistent with the activity value described by the manufacturer using the clot-based Limulus amebocyte lysate (LAL) assay E-Toxate test (Sigma-Aldrich) or using the microplate chromogenic LAL assay (LAL Endochrome-K, Endosafe; Charles River). Diphosphoryl Lipid A purified from *E. coli* F583 (Rd mutant) was purchased from Sigma Aldrich (L5399). The manufacturer specification for Lipid A endotoxin activity was 1 x 10^6^ EU per mg. Using a set of standard LPS dilutions, our results were lower than the activity described by the manufacturer, but Lipid A activity was consistently replicated using either the E-Toxate clot-based LAL or the Endochrome-K, Endosafe chromogenic LAL assay.

### Limulus amebocyte lysate gel assay

Detection and semi-quantitation of the endotoxin activities of LPS and Lipid A were performed using the tube-based E-Toxate test (Sigma-Aldrich). The LAL reagent, prepared from a lysate of circulating amebocytes of the horseshoe crab *Limulus polyphemus*, changes viscosity and opacity on contact with minute quantities of endotoxin (18). Endotoxin in the presence of calcium ions activates a trypsin-like enzyme that proteolytically modifies a “coagulogen” to produce clotted protein (19). The limit of sensitivity of the test is 0.05-0.10 endotoxin units (EU)/mL. The LAL measured activity is proportional to the pathophysiologic activity of LPS (20).

### Gel LAL inhibition testing

Detoxification of endotoxin by exposure to sodium hydroxide (NaOH) can be quantified using the E-Toxate test to measure *Limulus* amebocyte gel activity (20). In similar manner, the inhibitory effects of MPO and EPO on lipopolysaccharide (LPS) was investigated by contacting varying quantities of MPO or EPO with a given quantity of LPS and measuring for inhibition of endotoxin activity using the LAL gel assay.

### Chromogenic Limulus amebocyte lysate assay

Endotoxin was also quantified using a microplate chromogenic Limulus amebocyte lysate (LAL) assay (LAL Endochrome-K, Endosafe) purchased from Charles River (21). Endotoxin activation of the LAL clotting enzyme was quantified by measuring endotoxin-activated enzymatic cleavage of a synthetic chromogenic substrate releasing p-nitroaniline (pNA). Activity was measured as change in absorbance at a wavelength of 405 nm in a microplate spectrophotometer (Tecan). Kinetic measurement of the time to maximum color change was used to gauge the activity of endotoxin present. The limit for detection was set at 1680 seconds (28 min). Calibration standards were prepared and a standard curve was used to generate an equation with a coefficient of determination (R^2^) for each experiment performed.

### Chromogenic LAL inhibition testing

The modified chromogenic LAL method served as an *in vitro* assay of haloperoxidase inhibition of endotoxin. This inhibition assay included a pre-incubation step where either LPS or Lipid A was contacted with the test enzyme (MPO, EPO, or GO) for a period of 30 minutes at 37˚C. Test agents were diluted in low endotoxin reagent water (LRW). Following incubation Limulus lysate solution was added and chromogenic activity was measured as the time (in seconds) to maximum color change. Inhibition was calculated as the difference between the activity expected for the quantity of endotoxin present in the absence of haloperoxidase, and the actual measured endotoxin activity of LPS or Lipid A contacted with MPO or EPO. Inhibition was expressed as a percentage of the activity of LPS or Lipid A alone. MS Excel software was used for data analysis, curve fitting, coefficient of determination (R^2^) calculations and graphic construction. The R^2^ statistic tests the fit of the measured observations in proportion to total variation of outcomes predicted using the empirically generated equation; i.e., the proportion of the variance in the dependent variable predictable from the independent variable (22).

### Mice lethal dose 90 (LD90) model

Experimentally naïve, healthy BALB/c female mice with a weight range of 16.2 to 19.7 g were divided into dose groups. Treatment of the animals (including but not limited to all husbandry, housing, environmental, and feeding conditions) was conducted in accordance with the guidelines recommended in Guide for the Care and Use of Laboratory Animals. All mouse testing was performed according to the protocols and standard operational procedures of Concord Bioscience LLC, Concord, OH 44077.

Except for one group containing 15 mice, each test group contained 20 mice. For all groups each mouse received a 0.5 mL total volume with different doses of purified LPS injected intraperitoneally (IP). Low endotoxin reagent water (LRW) was used to adjust the concentrations of LPS. Groups 1, 2, 3 and 4 received 0.20, 0.35, 0.50 and 0.65 mg of LPS per mouse, respectively. After 5 days of observation, the censored (live) and the event (dead) mouse counts were tabulated. Group 1 had 20 live with 5 dead for a 75% survival (25% mortality); Group 2 had 5 live with 15 dead for a 25% survival (75%mortality), and Groups 3 and 4 had 20 dead with 0 live for 0% survival (100% mortality). The 90% lethal dose (LD90), i.e., the dose estimated to produce 90% mortality, was set at 0.40 mg per mouse, i.e., about 22 mg/kg.

The mouse LD90 testing was done in two parts. The first part tested the inhibitory action of 0.5 mg MPO combined with the 0.4 mg LPS (LD90 dose). The second part expanded the testing to include doses of 2.5 and 5.0 mg MPO, as well as doses of 2.5 and 5.0 mg EPO in combination with the 0.4 mg LPS (LD90 dose). The appropriate concentrations of LPS plus MPO or EPO were vortexed vigorously for about a minute then incubated at 37°C for 45 min. The mix was vortexed again prior to IP injection of each animal.

Survivor analysis was performed with IBM SPSS software using the Laerd Statistics guide for Kaplan Meier survival analysis.

## Results

### MPO and EPO inhibit LPS endotoxin activity measured by the clot-endpoint LAL

Endotoxin was quantified using the E-Toxate clot-endpoint Limulus amebocyte lysate (LAL) assay. Following testing of LPS standards and confirmation of endotoxin unit activity, LPS at 0.0125 EU per tube was tested in combination with a set of MPO and EPO dilutions to determine the lowest haloperoxidase dilution sufficient to completely inhibit clot-formation.

The results presented in **Table 1** demonstrate that both MPO and EPO inhibit the EU activity of LPS measured by the E-Toxate LAL gelation test. Based on the haloperoxidase mass required for inhibition, MPO is about 3.2 times more potent than EPO.

**TABLE 1.** Haloperoxidase inhibition of endotoxin activity (EU) activity measured as Limulus amebocyte lysate (LAL) gelation.

| LPS | EU/tube | None | 0.0250 | 0.0125 | 0.0060 | 0.0030 | 0.0015 |
| --- | --- | --- | --- | --- | --- | --- | --- |
|  |  | Neg | Pos | Pos | Pos | Neg | Neg |

Haloperoxidase inhibition of EU activity measured as LAL gelation
| LPS | EU/tube | None | 0.0125 |  |  |  |  |  |  |  |  |
| --- | --- | --- | --- | --- | --- | --- | --- | --- | --- | --- | --- |
| MPO | mg/tube | 0.560 | 0.560 | 0.420 | 0.320 | 0.240 | 0.180 | 0.130 | 0.100 | 0.075 | 0.056 |
|  |  | Neg | Neg | Neg | Neg | Neg | Neg | Neg | Neg | Pos | Pos |
| EPO | mg/tube | 0.560 | 0.560 | 0.420 | 0.320 | 0.240 | 0.180 | 0.130 | 0.100 | 0.075 | 0.056 |
|  |  | Neg | Neg | Neg | Neg | Pos | Pos | Pos | Pos | Pos | Pos |

### Chromogenic LAL assay measurement of MPO and EPO inhibition of LPS

Endotoxin was also quantified using the chromogenic Limulus amebocyte lysate (LAL) assay (21). Endotoxin activated cleavage of the chromogenic substrate releases p-nitroaniline that is measured at a wavelength of 405 nm with microplate spectrophotometer. Temporal kinetic measurement of the time-to-maximum-slope was used to gauge the endotoxin activity.

Calibration standards were run and a standard curve with equation was generated for each set of measurements as shown in Fig. 1 Graph A.

**Fig. 1.**
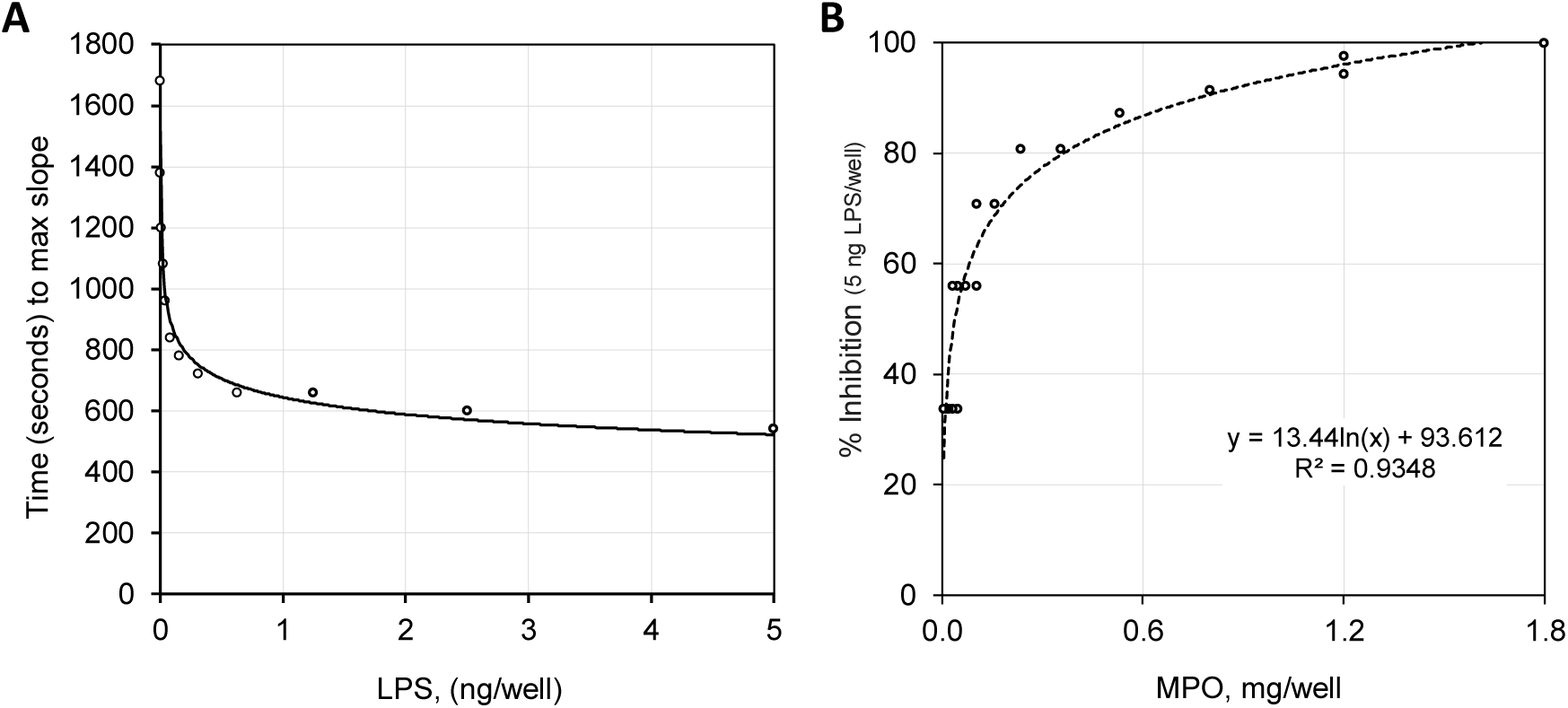
**Graph A.** Regression plot of LPS standards (ng/well) against time (sec) to maximum slope. For the range: 0.0012 - 5 ng LPS/well the derived equation was y = 643.96x^-0.131^ with an R² = 0.982; for the range: 0.0012 - 0.625 ng LPS/well the equation was y = 600.99x^-0.146^ with an R² = 0.991. **Graph B.** Percent inhibition of the LAL activity for 5.0 ng LPS/well plotted against varying concentrations of MPO (mg/well). Testing was run in quadruplicate with all data points depicted. The regression equation shows inhibition proportional to the natural log of the MPO mass with R^2^ is included.

Using a set of standards, the endotoxin mass/well is plotted against the time point of maximum color change (max slope). The manufacturer specified activity for this lot of LPS was 3 x 10^6^ endotoxin units (EU) per mg, i.e., 3 EU/ng. As expected, the 0.01 ng LPS/well standard had a time to max slope of 1200 sec, a value equivalent to about 0.03 EU. This chromogenic LAL test was modified to serve as an *in vitro* assay of haloperoxidase inhibition of endotoxin. The inhibition assay included a pre-incubation step where LPS or Lipid A was exposed to a test enzyme (MPO, EPO, or GO) for a period of 30 minutes at 37˚C. The reaction conditions contained no hydrogen peroxide, and as such, inhibition reflected protein binding in the absence of haloperoxidase enzymatic action. Following incubation, reaction was initiated by addition of the Limulus lysate solution, and the time (in seconds) to maximum color change was measured. Based on the standards-derived equation, the time-to-max-slope was converted to equivalent LPS mass. Inhibition was calculated as the difference between the activity of the expected mass, e.g., endotoxin without MPO, and the actual measured LPS activity, e.g., the percentage of endotoxin activity measured in the presence of MPO. The plot of MPO mass versus % of LPS endotoxin activity is illustrated by the plot of the data shown in **Fig. 1 Graph B**.

Inhibition of the endotoxin activity of 5.0 ng LPS was proportional to the log of MPO concentration (mass) present. One mg MPO inhibited about 91% of the endotoxin activity of 5 ng LPS; stated differently, 1 mg MPO inhibited about 4.5 ng LPS. However, at one-hundredth the MPO mass (0.01 mg), MPO was still capable of inhibiting 28% of the activity of 5 ng LPS, i.e., about 1.4 ng LPS. Note that with LPS mass held constant and MPO mass varied, the percent inhibition of LAL activity is directly proportional to the natural log of MPO concentration. As the ratio of the MPO to LPS mass increases, the percent inhibition of endotoxin activity increases, but the efficiency of MPO inhibition decreases.

For comparison, the endotoxin inhibition activities of MPO and EPO were measured and the results are plotted in **Fig. 2**. Both MPO and EPO inhibited LPS, but MPO was superior to EPO in inhibition capacity on a mass or molar basis. For a constant mass of LPS, inhibition of endotoxin activity was proportional to the log of the mass of MPO or EPO. At a mass of 0.25 mg, MPO and EPO inhibited 88% and 45% of the endotoxin activity of 0.1 ng LPS, respectively. Glucose oxidase (GO) did not inhibit LPS endotoxin activity within the range of concentration tested.

**Fig. 2.**
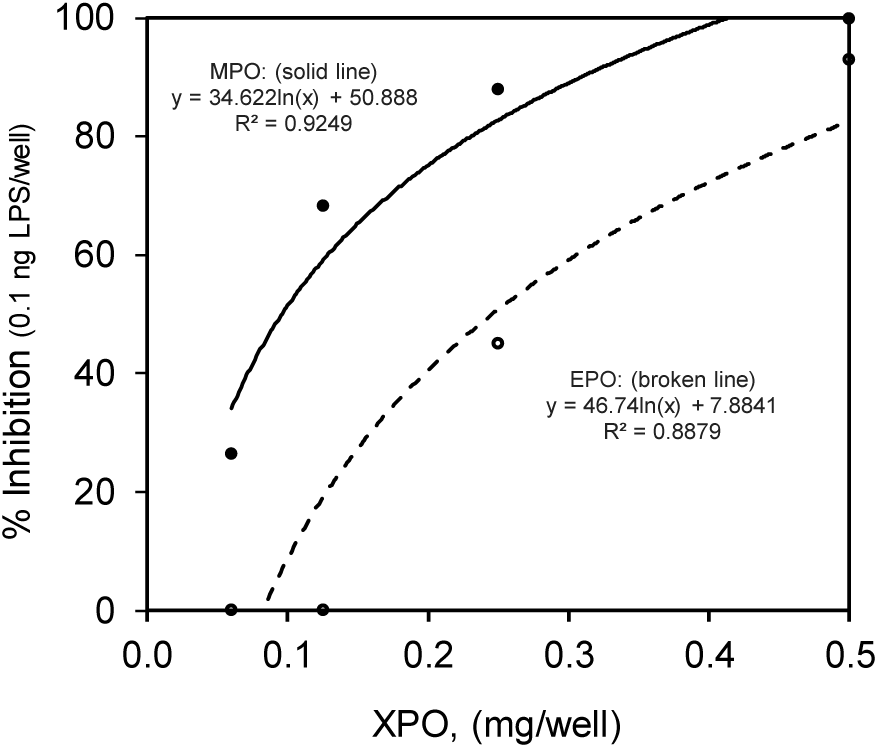
Percent inhibition of LAL endotoxin activity for 0.1 ng LPS/well plotted against mg/well of MPO, EPO and GO (not shown). The regression equations are shown for MPO and for EPO. LPS inhibition is in proportion to the natural log of the MPO and EPO mass contacted. GO was not inhibitory and not included.

### MPO and EPO inhibition of Lipid A endotoxin activity by the chromogenic LAL

Lipid A, the toxic component of LPS (14), has two glucosamine units with anionic phosphate groups attached and typically six hydrophobic fatty acids that anchor it into the outer membrane of GNB. Purified Lipid A is toxic and its endotoxin activity is measurable with the LAL assay.

The inhibitory actions of MPO and EPO were tested on diphosphoryl Lipid A from *E. coli* F583 (Sigma Aldrich, L5399) using the same chromogenic LAL assay described above. The results of Lipid A standards testing over time are presented in **Fig. 3 Graph A**.

**Fig. 3.**
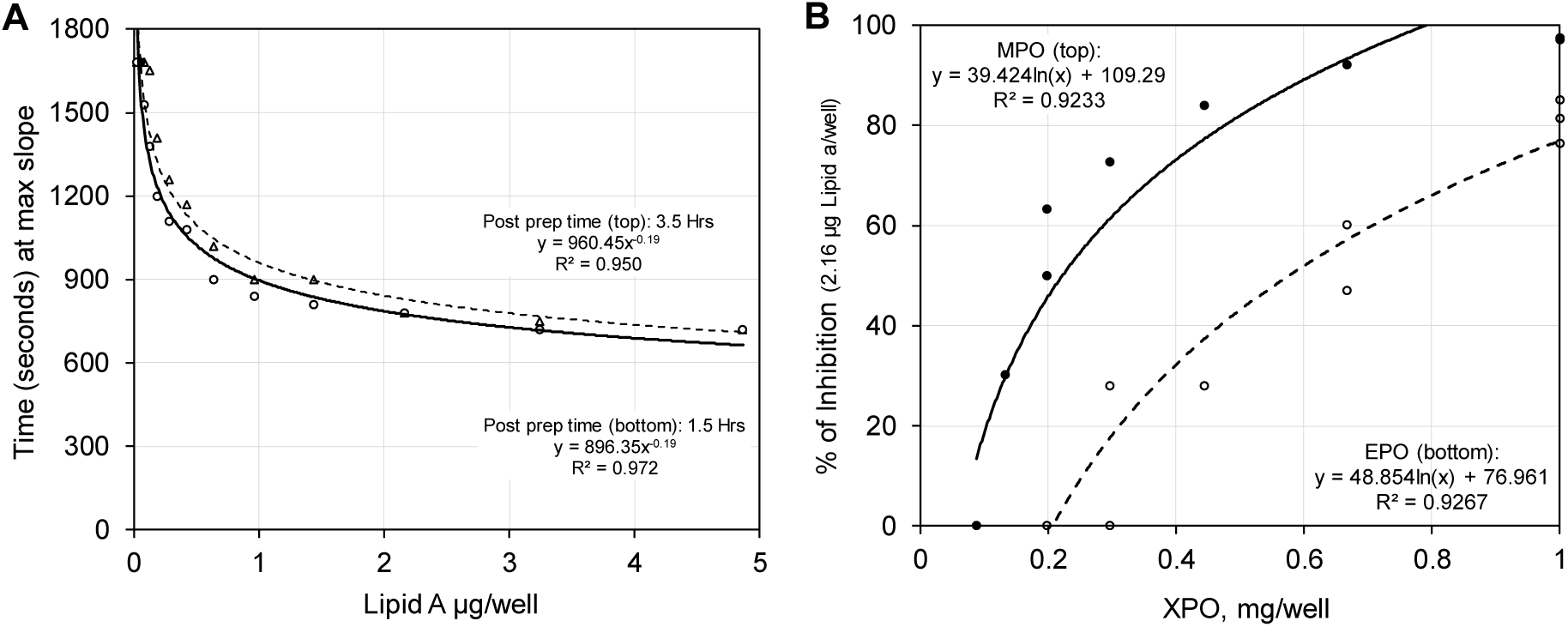
**Graph A.** Regression plot of Lipid A standards (ng/well) against time (sec) to maximum slope with derived equations and R^2^’s. The top and bottom curves were obtained from the same set of standards prepared 1.5 and 3.5 hours post preparation of the Lipid A standards as indicated. **Graph B.** Percent inhibition of LAL activity for 2.16 µg Lipid A/well plotted against varying concentrations of MPO or EPO in mg/well. The regression equations for MPO and EPO inhibition of Lipid A are shown. Lipid A standards were simultaneously run for each MPO or EPO inhibition experiment.

The hydrophobic character of Lipid A complicated preparation of aqueous standards (13), and the endotoxin unit activity per Lipid A mass was lower than the manufacture reported value (i.e., ≥ 1 × 10^6^ EU/mg) based a coagulation LAL assay. As illustrated by the data of **Fig. 3 Graph A**, Lipid A endotoxin activity quantified using the chromogenic LAL assay was reasonably stable over several hours. Using the equation generated from simultaneously run Lipid A standards, MPO and EPO inhibition of Lipid A endotoxin activity was measured as shown in **Fig. 3 Graph B**. Consistent with the finding for LPS, the endotoxin activity of Lipid A was inhibited by both MPO and EPO, and inhibition was proportional to the natural log of the MPO or EPO concentration. Likewise, MPO was more inhibitory than EPO. GO did not inhibit the endotoxin activity of Lipid A.

### Haloperoxidase enzymatic action not required for endotoxin inhibition

The microbicidal and antitoxin activities of MPO require enzymatic action, i.e., haloperoxidase activity (6, 12, 23). All of the experiments described above were conducted in the absence of hydrogen peroxide or a peroxide generating system, i.e., in the absence of haloperoxidase enzymatic action. This experiment was devised to measure possible differences in endotoxin inhibition with regard to haloperoxidase activity. A formulation containing MPO plus GO as a hydrogen peroxide generating enzyme, i.e., 4 µg MPO and 0.8 µg GO. The enzyme complex was tested for LPS endotoxin inhibitory activity in the absence of D-glucose, i.e., no haloperoxidase action, and in the presence of D-glucose, i.e., high haloperoxidase activity. The microbicidal action of the fully active MPO-GO-glucose preparation has been previously described (10, 24). The results are presented in **Fig. 4**.

**Fig. 4.**
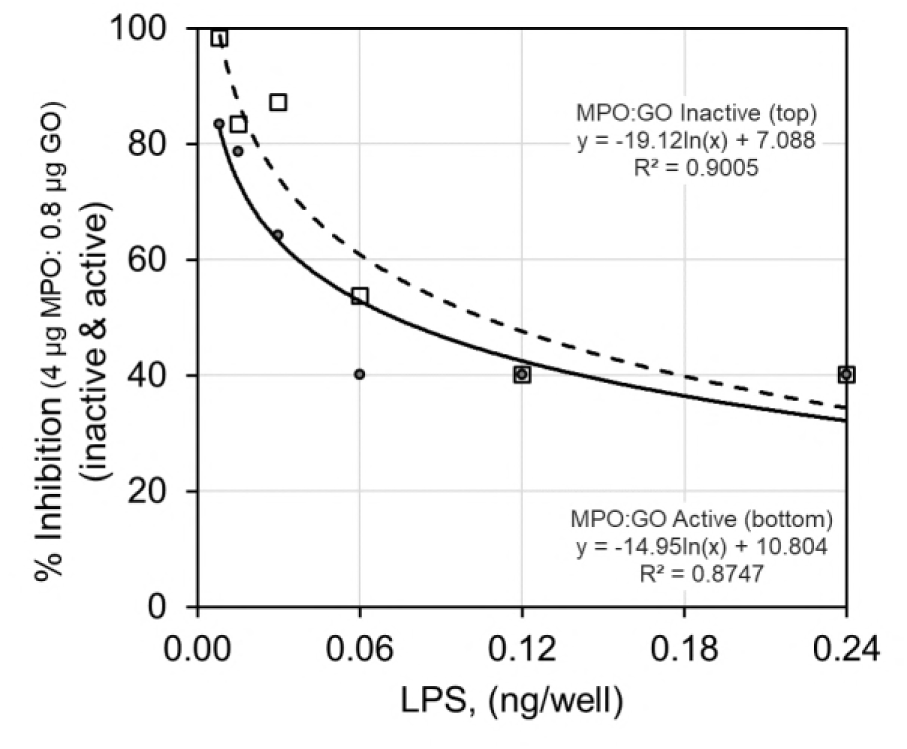
Percent inhibition of LPS endotoxin activity for a formulation composed of 4 µg MPO and 0.8 µg GO with 23 µg Cl^-^/well measured against varying concentrations of LPS using the LAL assay. The formulation was tested without D-glucose, i.e., no haloperoxidase activity, and with a non-limiting concentration (0.2 mg/well) of D-glucose, i.e., haloperoxidase activity. The regression equations describe that the percentage inhibition of LPS endotoxin activity for a constant concentration of MPO plus GO with or without D-glucose is proportional to the negative log of the LPS concentration.

With the concentration of MPO plus GO held constant, i.e., MPO held constant, and the concentration of LPS varied, the percent inhibition of endotoxin activity is proportional to the negative natural log of the LPS concentration. This is the expected inverse relationship to that observed when LPS or Lipid A is held constant and MPO or EPO is varied. As the mass ratio of LPS to MPO increases, the percent inhibition of endotoxin activity decreases. The functional enzymatic MPO-GO complex, with a non-limiting concentration of D-glucose as substrate and chloride as cofactor, was slightly less effective than the non-enzymatically active MPO-GO complex without D-glucose. Contacting MPO with LPS inhibited its endotoxin activity in the absence or presence of oxidative enzymatic function. Haloperoxidase activity did not improve endotoxin inhibition.

### Effect of MPO and EPO on mouse survival in an endotoxin LD90 model

Intraperitoneal (IP) injection of a 0.5 mL volume containing 0.4 mg LPS per mouse produced about 90% lethality (LD90) in the naïve, healthy BALB/c female mice tested as described in **Methods**. The LD90 dose, equivalent to about 22 mg/kg, was used for all testing. Highly purified porcine MPO and EPO were used for testing as described in **Materials**. The indicated concentrations of LPS and MPO were mixed for about a minute, incubated at 37°C for 45 min, then vortexed again prior to intraperitoneal injection (IP) of 0.5 mL per mouse. No hydrogen peroxide was present in the preparation, and as such, there was MPO contact without haloperoxidase action.

LD90 testing consisted of two studies. The initial study determined the effect of 0.5 mg of MPO on the survivability of mice treated with 0.4 mg of LPS. As depicted in **Figure 5 A** and described in **Table 2**, this quantity of MPO significantly improved the survival of mice in the LPS LD90 model (25-29). Contacting the LD90 dose of 0.4 mg LPS with 0.5 mg MPO resulted in a 50% survival rate over the expected 10% survival rate. MPO-treatment did not appear to change the incidence of LPS-associated clinical observations, but LPS-associated mortality was clearly and significantly decreased in the MPO-treated animals, as depicted in the Kaplan Meier plot of **Fig. 5 Graph A.**

**Fig. 5.**
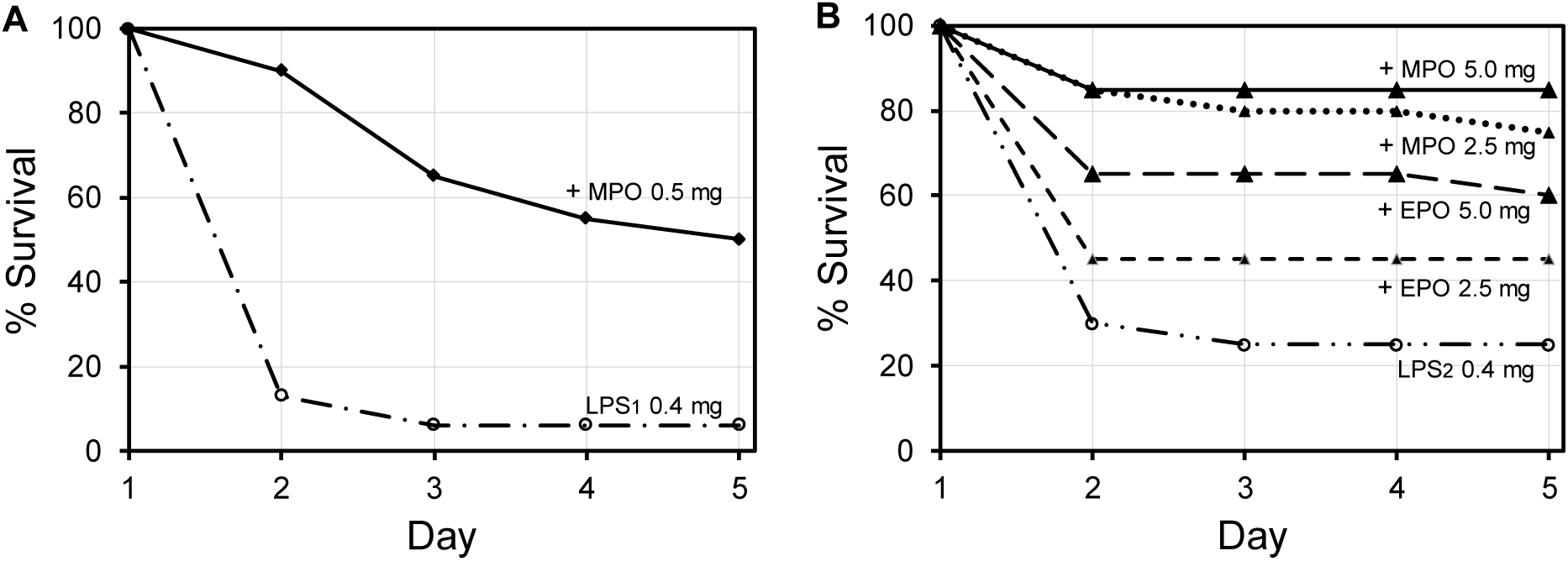
**Graph A.** Initial study of the effect of 0.5 mg MPO/mouse on survival depicted as Kaplan Meier survivor curves for BALB/c female mice over a 5-day period following IP injection of an LD90 (0.4 mg LPS/mouse; i.e., about 22 mg/kg) dose on Day 1. **Graph B.** Follow-up study showing Kaplan Meier survivor curves using 2.5 and 5.0 mg MPO/mouse, and also, 2.5 and 5.0 mg EPO/mouse.

**TABLE 2.**
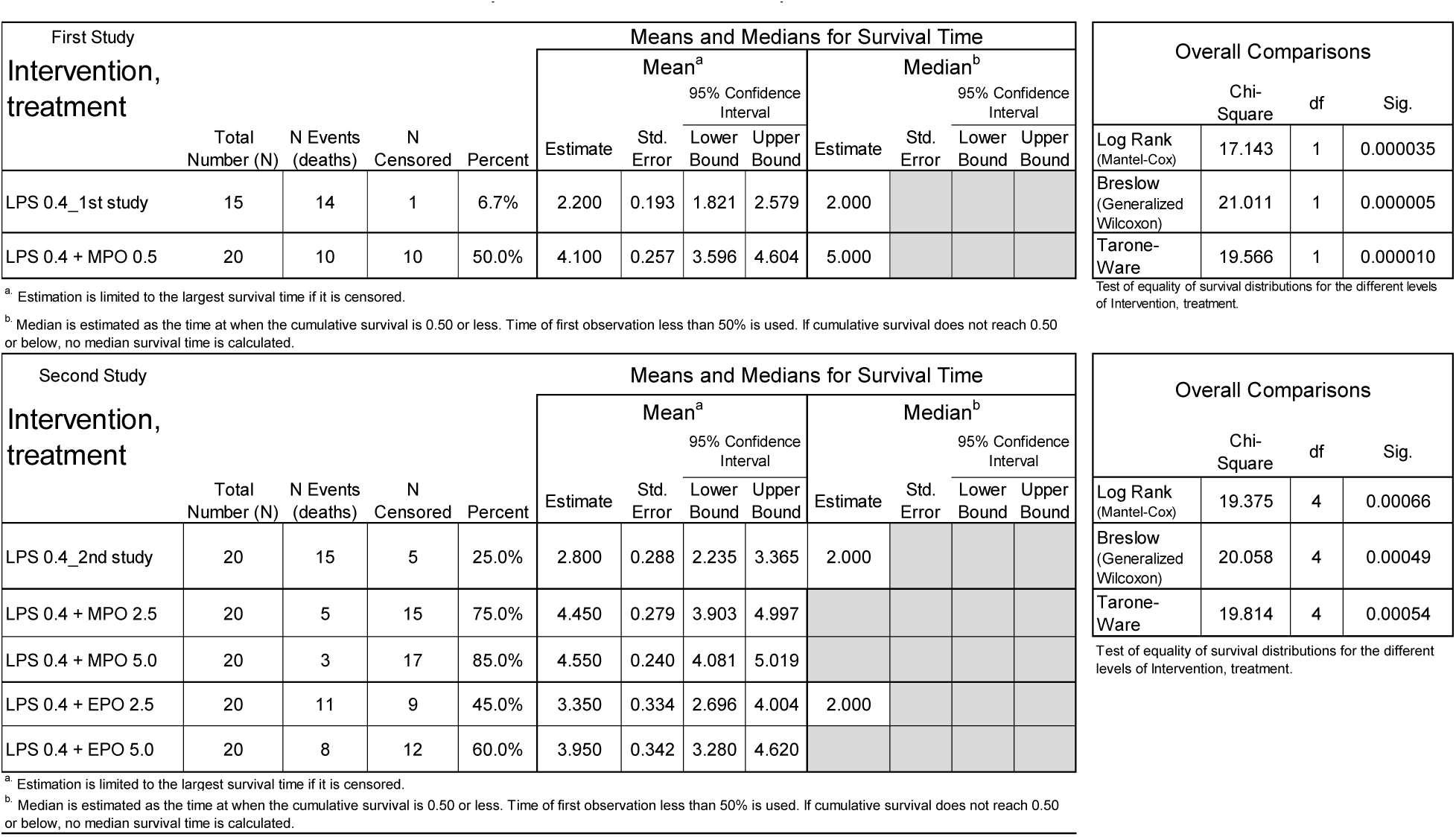
Mean and median survival times plus overall statistical comparisons for the first and second LD90 mouse studies.

A second LD90 study was designed to confirm the initial results, expand the concentration range of MPO, and also include the same range of EPO for comparison. The LPS only LD90 group for the second study showed less than the expected mortality, i.e., 75% mortality. Overall comparison of the LD90 controls for the initial and second studies using Log Rank (Mantel-Cox) analysis yields a chi-square of 2.248 with 1 degree of freedom (df) for a significance (p value) of 0.134, i.e., no significant difference in the LD90 controls (27). As an additional control, a group of twenty mice were treated with high dose (5 mg/mouse) MPO without LPS. The mice treated with this high dose of MPO alone (without LPS) showed no abnormal clinical observations and no mortality. However, an EPO only control group of twenty mice treated with high dose (5 mg/mouse) EPO without LPS showed 10% lethality by day 2. The surviving EPO only group mice showed no abnormal clinical observations.

As depicted in **Figure 5 B** and described in the overall analysis of **Table 2** and in the pairwise analysis of **Table 3**, the increasing the concentrations of MPO to 2.5 and 5.0 mg/mouse yielded significantly improved mouse survival in this LPS LD90 model, and survival was increased in a dose dependent manner. The EPO group tested at concentrations of 2.5 and 5.0 mg/mouse also showed improved mouse survival in this LPS LD90 model, and survival was increased in a dose dependent manner as depicted in **Figure 5 B**. However, the small increase in survival for the 2.5 mg EPO/mouse group was not significant; the pairwise Kaplan Meier analysis of **Table 3** showed a p value of 0.1965 (27). At the higher EPO concentration of 5.0 mg/mouse, a statistically significant increase in mouse survival was obtained, i.e., p value of 0.0235.

**TABLE 3.** Pairwise comparison of LPS alone and in combination with different concentrations of MPO and EPO.

| Intervention, treatment |  | Pairwise Comparisons |  |  |  |  |  |  |  |
| --- | --- | --- | --- | --- | --- | --- | --- | --- | --- |
|  |  | LPS 0.4 + MPO 2.5 |  | LPS 0.4 + MPO 5.0 |  | LPS 0.4 + EPO 2.5 |  | LPS 0.4 + EPO 5.0 |  |
|  |  | Chi-Square | Sig. | Chi-Square | Sig. | Chi-Square | Sig. | Chi-Square | Sig. |
| Log Rank (Mantel-Cox) | LPS 0.4 (2nd study) | 10.782 | 0.0010 | 14.421 | 0.0001 | 1.668 | 0.1965 | 5.131 | 0.0235 |

## Discussion

MPO binds to a broad spectrum of microbes. Strong binding has been observed with many Gram-positive bacteria (GPB), including *Staphylococcus aureus*, and with all Gram-negative bacteria (GNB) tested. Members of the lactic acid family of GPB show relatively weak binding (30-32). The strength of MPO binding is proportional to the efficiency of haloperoxidase-mediated microbicidal action. When the concentration of MPO is limiting, strong MPO-binding to bacteria such as *S. aureus*, *Escherichia coli*, or *Pseudomonas aeruginosa* in combination with weak MPO-binding to hydrogen peroxide-producing viridans streptococci results in restriction of microbicidal action to MPO-bound bacteria and sparing of streptococci. This selectivity is observed in the absence and in the presence of strongly catalase-positive human erythrocytes at a high ratio of erythrocytes to MPO-binding bacteria, further supporting the requirement of proximity with regard to MPO microbicidal action. The primary and secondary microbicidal products of MPO action, i.e., hypochlorite and singlet molecular oxygen with its microsecond lifetime, favor the selective killing of MPO-bound microbes (10, 32).

MPO inactivates microbial toxins including diphtheria toxin, tetanus toxin, and *Clostridium difficile* cytotoxin (11, 12, 23). The toxin-destroying activities of MPO required haloperoxidase enzymatic action. Enzymatic or non-enzymatic action of MPO or EPO on endotoxin by MPO or EPO has not been described. The moderate to strong MPO binding reported for all GNB might be related to the fact that endotoxin is a characteristic of GNB (14-16). All GNB tested to date show MPO binding (10, 31), suggesting a possible commonality with regard to the mechanism of binding. The cell envelope of a GNB is composed of a cell wall with an inner cytoplasmic cell membrane and an outer cell wall membrane presenting lipopolysaccharide (LPS) with its toxic Lipid A component (13). The toxic action of LPS remains even after the death of the GNB. Release of endotoxin secondary to GNB lysis produces severe toxemia, i.e., septic shock.

The present studies were designed to determine if haloperoxidase binding, in addition to improving the potency and selectivity of microbicidal action, might provide additional protection against the endotoxin activity of residual LPS and Lipid A. The presented results demonstrate that MPO, and to a lesser extent EPO, bind to and inhibit the endotoxin activity of LPS measured by either the gel or chromogenic LAL assay. Electrostatic interaction is reasonably expected to play a role in MPO and EPO binding to the anionic phosphate groups of LPS and its Lipid A component. Electrostatic binding alone is insufficient to explain the threefold greater inhibitory action of MPO relative to more cationic EPO. MPO is significantly less cationic than EPO. Although the mechanism of inhibition likely involves electrostatic interaction, electrostatic binding provides an incomplete explanation.

The toxic activities of LPS and Lipid A are measured in endotoxin units (EU). Measuring EU activity using the chromogenic LAL assay, the percent inhibition of a constant concentration of LPS or Lipid A is directly proportional to the natural log of the MPO or EPO concentration as shown in **Figures 1-3**. Likewise, if the concentration of MPO is held constant and the LPS concentration is varied, the percent inhibition of endotoxin activity is proportion to the negative natural log of the LPS concentration as shown in **Figure 4**.

MPO or EPO binding is sufficient for inhibition of LPS and Lipid A endotoxin activity. Unless stated otherwise, no hydrogen peroxide was available to either MPO or EPO. As described in Figure 4, when endotoxin inhibition was tested in the absence and in the presence of a hydrogen peroxide generation, the enzyme-functional haloperoxidase system produced slight decreased inhibition. MPO and EPO haloperoxidase enzymatic function are not required for endotoxin inhibition.

Although the clot-based and chromogenic assays for MPO and EPO inhibition of endotoxin activities provide good *in vitro* evidence, we extend testing to include an *in vivo* LD90 mouse pathophysiology model of LPS endotoxicity. Initial study showed strong evidence for increased survivability using MPO at 0.5 mg relative to the LD90 dose of 0.4 mg LPS. The follow-up study increased the doses to 2.5 and 5.0 mg MPO and expanded testing to include testing of 2.5 and 5.0 mg EPO; the same LD90 dose of 0.4 mg LPS was used. As depicted by the Kaplan Meier plots of **Figure 5**, MPO, and to a lesser extent EPO, increased survivability in in a dose dependent manner. As indicated by statistical analysis of the results presented in Tables 2 and 3, increased survival was statistically significant at all MPO concentrations tested. However, statistically significant increase in survival was only achieved using 5.0 mg EPO.

The *in vitro* and *in vivo* results presented herein unambiguously demonstrate that MPO, and to a lesser extent EPO, inhibit endotoxin activity. Thus, in addition to the potent haloperoxidase microbicidal action of MPO and EPO, these proteins also provide non-enzymatic protection against endotoxin, and this protection is independent of microbicidal enzymatic action. Our position is that the haloperoxidase action of MPO is pH-restricted, and that the selective binding characteristic of MPO physically focuses its potent but temporally transient oxygenation activities. The enzymatic action of MPO is focused to exert maximum microbicidal action and produces minimal host damage (10). The direct *in vitro* and *in vivo* evidence of MPO binding inhibition of endotoxin activity presented herein is consistent with the recently reported evidence that blockade or genetic deletion of MPO increases the mortality associated with LPS toxicity, and that MPO protects against the adverse effects of endotoxin (17).

